# Changes in temperature alter susceptibility to a virus following a host shift

**DOI:** 10.1101/358564

**Authors:** Katherine E. Roberts, Jarrod D. Hadfield, Manmohan D. Sharma, Ben Longdon

## Abstract

Host shifts - where a pathogen jumps between different host species - are an important source of emerging infectious disease. With ongoing climate change there is an increasing need to understand the effect changes in temperature may have on emerging infectious disease. We investigated whether species’ susceptibilities change with temperature and ask if susceptibility is greatest at different temperatures in different species. We infected 45 species of *Drosophilidae* with an RNA virus and measured how viral load changes with temperature. We found the host phylogeny explained a large proportion of the variation in viral load at each temperature, with strong phylogenetic correlations between viral loads across temperature. The variance in viral load increased with temperature, whilst the mean viral load did not, such that as temperature increased the most susceptible species become more susceptible, and the least susceptible less so. We found no significant relationship between a species’ susceptibility across temperatures and proxies for thermal optima; critical thermal maximum and minimum or basal metabolic rate. These results suggest that whilst the rank order of species susceptibilities can remain the same with changes in temperature, the likelihood of host shifts into a given species may increase or decrease.

**Author Summary:** Emerging infectious diseases are often the result of a host shift, where a pathogen jumps from one host species into another. Understanding the factors underlying host shifts is a major goal for infectious disease researchers. This effort has been further complicated by the fact that host-parasite interactions are now taking place in a period of unprecedented global climatic warming. Here, we ask how host shifts are affected by temperature by carrying out experimental infections using an RNA virus across a wide range of related species, at three different temperatures. We find that as temperature increases the most susceptible species become more susceptible, and the least susceptible less so. This has important consequences for our understanding of host shift events in a changing climate, and suggests that temperature changes may affect the likelihood of a host shift into certain species.

## Introduction

Temperature is arguably the most important abiotic factor that affects all organisms, having both indirect and direct effects on physiology and life history traits [1–3]. There is much to be learned about the impact of climate change on infectious diseases [1,4,5]. Changes in temperature can impact both host and parasite biology, leading to complex and difficult to predict outcomes [2,6].

Host shifts, where a parasite from one host species invades and establishes in a novel host species, are an important source of emerging infectious disease [7]. Some of the most deadly outbreaks of infectious diseases in humans including Ebola virus, HIV and SARS coronavirus have been linked to a host switch event [8–11] and many others have direct animal vectors or reservoirs (e.g. Dengue and Chikungunya viruses) [12,13]. The potential for novel host shifts may increase with changing temperatures due to fluctuations in host and/or parasite fitness, or changes in species distributions and abundances [14,15]. Distribution changes may lead to new species assemblages, causing novel contacts between parasites and potential hosts [16–18].

Susceptibility to infection is known to vary with temperature due to within individual physiological changes in factors such as the host immune response, metabolic rate or behavioural adaptations [19–22]. Thermally stressed hosts may face a trade-off between the resource investment needed to launch an immune response versus that needed for thermoregulation, or behavioural adaptations to withstand sub-optimal temperatures [23–26]. Temperature shifts could also cause asymmetrical or divergent effects on host and parasite traits [27]. For example, changes in temperature may allow differential production and survival of parasite transmission stages, and changes in replication rates, generation times, infectivity and virulence [28–30].

Host shifts have been shown to be more likely to occur between closely related species [31–33], but independently of this distance effect, clades of closely related hosts show similar levels of susceptibility [34,35]. Thermal tolerances - like virus susceptibility - are known to vary across species, with groups of closely related species having similar thermal limits, with a large proportion of the variation in these traits being explained by the phylogeny [36–39]. Previous studies on host shifts have assayed the susceptibility of species at a single temperature [32,34,35,40]. However, if the host phylogeny also explains much of the variation in thermal tolerance, then phylogenetic patterns in virus susceptibility could be due to differences between species’ optima and the chosen assay temperatures. Therefore, for experiments carried out at a single temperature, phylogenetic signal in thermal tolerance may translate into phylogenetic signal in thermal stress. Any apparent phylogenetic signal in susceptibility could potentially be due to the effects of thermal stress, and may not hold true if each species was to be assayed at its optimal temperature.

Here, we have asked how species susceptibilities change at different temperatures and whether susceptibility is greatest at different temperatures in different species. We infected 45 species of *Drosophilidae* with Drosophila C Virus (DCV; Dicistroviridae) at three different temperatures and measured how viral load changes with temperature. We also examine how proxies for thermal optima and cellular function (thermal tolerances and basal metabolic rate) relate to virus susceptibility across temperatures [37–39]. DCV is a positive sense RNA virus in the family *Discistroviridae* that was isolated from *Drosophila melanogaster* and naturally infects several species of Drosophilidae in the wild [41–43]. DCV infected flies show reduced metabolic rate and activity levels, develop an intestinal obstruction, reduced hemolymph pH and decreased survival [44–47]. This work examines how temperature can influence the outcomes of host shifts, and looks at some of the potential underlying causes.

## Methods

### Experimental infections

We used Drosophila C virus (DCV) clone B6A, which is derived from an isolate collected in Charolles, France [48]. The virus was prepared as described previously [49]; briefly DCV was grown in Schneider’s Drosophila line 2 cells and the Tissue Culture Infective Dose 50 (TCID**50**) per ml was calculated using the Reed-Muench end-point method [50].

Flies were obtained from laboratory stocks of 45 different species. All stocks were maintained in multi generation populations, in Drosophila stock bottles (Dutscher Scientific) on 50ml of their respective food medium at 22°C and 70% relative humidity with a 12 hour light-dark cycle (see table S1 for rearing conditions for each species). Each day, two vials of 0-1 day old male flies were randomly assigned to one of three potential temperature regimes; low, medium or high (17°C, 22°C and 27 °C respectively) at 70% relative humidity. Flies were tipped onto fresh vials of food after 3 days, and after 5 days of acclimatisation at the experimental temperature were infected with DCV. Flies were anesthetized on CO_2_ and inoculated using a 0.0125 mm diameter stainless steel needle that was bent to a right angle ~0.25mm from the end (Fine Science Tools, CA, USA). The bent tip of the needle was dipped into the DCV solution (TCID**50** = 6.32×10^9^) and pricked into the pleural suture on the thorax of the flies. One vial of inoculated flies was immediately snap frozen in liquid nitrogen to provide a time point zero sample as a reference to control for relative viral dose. The second vial of flies were placed onto a new vial of fresh cornmeal food and returned to their experimental temperature. After 2 days (+/− 1 hour) flies were snap frozen in liquid nitrogen, this time point was chosen based on pilot data as infected flies showed little mortality at 2 days post infection, and viral load plateaus from day 2 at 22°C. Temperatures were rotated across incubators in each block to control for incubator effects. All frozen flies were homogenised in a bead homogeniser for 30 seconds (Bead Ruptor 24; Omni international, Georgia, USA) in Trizol reagent (Invitrogen) and stored at - 80°C for later RNA extractions.

These collections and inoculations were carried out over three replicate blocks, with each block being completed over consecutive days. The order that the fly species were infected was randomized each day. We aimed for each block to contain a day 0 and day 2 replicate for each species, at each temperature treatment (45 species × 3 temperatures × 3 experimental blocks). In total we quantified viral load in 12,827 flies, with a mean of 17.1 flies per replicate (range across species = 4-27). Of the 45 species, 44 had 6 biological replicates and one species had 5 biological replicates.

### Measuring the change in viral load

The change in RNA viral load was measured using qRT-PCR. Total RNA was extracted from the Trizol homogenised flies, reverse-transcribed with Promega GoScript reverse transcriptase (Promega) and random hexamer primers. Viral RNA load was expressed relative to the endogenous control housekeeping gene *RpL32 (RP49). RpL32* primers were designed to match the homologous sequence in each species and crossed an intron-exon boundary so will only amplify mRNA [34]. The primers in *D. melanogaster* were *RpL32* qRT-PCR F (5’- TGCTAAGCTGTCGCACAAATGG -3’) and *RpL32* qRT-PCR R (5’- TGCGCTTGTTCGATCCGTAAC -3’). DCV primers were 599F (5’- GACACTGCCTTTGATTAG-3’) and 733R (5’CCCTCTGGGAACTAAATG-3’) as previously described [35]. Two qRT-PCR reactions (technical replicates) were carried out per sample with both the viral and endogenous control primers, with replicates distributed across plates in a randomised block design.

qRT-PCR was performed on an Applied Biosystems StepOnePlus system using Sensifast Hi-Rox Sybr kit (Bioline) with the following PCR cycle: 95°C for 2min followed by 40 cycles of: 95°C for 5 sec followed by 60°C for 30 sec. Each qRT-PCR plate contained four standard samples. A linear model was used to correct the cycle threshold (Ct) values for differences between qRT-PCR plates. Any samples where the two technical replicates had cycle threshold (Ct) values more than 2 cycles apart after the plate correction were repeated. To estimate the change in viral load, we first calculated *ΔCt* as the difference between the cycle thresholds of the DCV qRT-PCR and the *RpL32* endogenous control. The viral load of day 2 flies relative to day 0 flies was then calculated as 2^−ΔΔCt^, where *ΔΔCt* = *ΔCt_dayo_ -ΔCt_day2_*, where *ΔCt*_*dayo*_ and *ΔCt*_*day2*_ are a pair of *ΔCt* values from a day 0 biological replicate and a day 2 biological replicate for a particular species. Calculating the change in viral load without the use of the endogenous control gene *(RpL32)* gave equivalent results (Spearman’s correlation between viral load calculated with and without endogenous control: *ρ*= 0.97, *P<* 0.005)

### Critical Thermal Maximum and Minimum Assays

We carried out two assays to measure the thermal tolerances of species; a cold resistance measure to determine critical thermal minimum (CT_min_) under gradual cooling, and a heat resistance measure through gradual heating to determine critical thermal maximum (CT_max_). 0-1 day old males were collected and placed onto fresh un-yeasted cornmeal food vials. Flies were kept for 5 days at 22°C and 70% relative humidity and tipped onto fresh food every 2 days. In both assays individual flies were placed in 4 ml glass vials (ST5012, Ampulla, UK) and exposed to temperature change through submersion in a liquid filled glass tank (see supplementary material and methods for description of apparatus). For CT_max_ the tank was filled with water and for CT_min_ a mixture of water and ethylene glycol (50:50 by volume) was used to prevent freezing and maintain a constant cooling gradient. Five biological replicates were carried out for each species for both CT_max_ and CT_min_. Temperature was controlled using a heated/cooled circulator (TXF200, Grant Instruments, Cambridgeshire, UK) submerged in the tank and set to change temperatures at a rate of 0.1 °C/min, always starting from 22°C (the rearing temperature for stock populations). Flies were monitored continually throughout the assay and the temperature of knock down was ascertained by a disturbance method, whereby a fly was scored as completely paralysed if on gentle tapping of the vial wall the fly did not move any of its body parts.

### Measuring Metabolic Rate

To examine how cellular function changes with temperature, we estimated the resting metabolic rate of each species at 17°C, 22°C and 27 °C. Following the same methods as the viral inoculation assay, groups of 10, 0-1 day old male flies from 44 species were acclimatised at the three experimental temperatures for 5 days *(D. pseudobscura* was excluded as not enough individuals could be obtained from stocks for sufficient replication). Every 2 days flies were tipped onto fresh vials of cornmeal food. This was repeated in three blocks in order to get three repeat measures of metabolic rate for each of the species, at each of the three experimental temperatures. Flies were collected in a randomly assigned order across the three blocks

Closed system respirometry was used to measure the rate of CO_2_ production (VCO_2_) as a proxy for metabolic rate [51]. Flies were held in 10ml^−3^ airtight plastic chambers constructed from Bev-A-Line V Tubing (Cole-Parmer Instrument Company, UK). All measures were carried out during the day inside a temperature controlled incubator, with constant light, that was set to each of the experimental temperatures that the flies had been acclimatised to. The set up followed that of Okada *et al.* (2011) [52]. Compressed air of a known concentration of oxygen and nitrogen (21% O2:79% N_2_) was scrubbed of any CO_2_ and water (with Ascarite II & Magnesium Perchlorate respectively) and pumped through a Sable Systems RM8 eight-channel multiplexer (Las Vegas, NV, USA) at 100 ml/min^−1^ (±1%) into the metabolic chambers housing the groups of 10 flies. The first chamber was left empty as a reference cell, to acquire a baseline reading for all subsequent chambers at the start and end of each set of runs, therefore seven groups of flies were assayed in each run. Air was flushed into each chamber for 2 minutes, before reading the previous chamber. Readings were taken every second for 10 minutes by feeding the exiting air through a LiCor LI-7000 infrared gas analyser (Lincoln, NE, USA). Carbon dioxide production was measured using a Sable Systems UI2 analog-digital interface for acquisition, connected to a computer running Sable Systems Expedata software (v1.8.2) [53]. The metabolic rate was calculated from the entire 10-minute recording period, by taking the CO_2_ reading of the ex-current gas from the chamber containing the flies and subtracting the CO_2_ measure of the incurrent gas entering the chamber. These values were also corrected for drift away from the baseline reading of the empty chamber. Volume of CO_2_ was calculated as VCO_2_ = FR (Fe CO_2_ - Fi CO_2_) / (1-Fi CO_2_). Where FR is the flow rate into the system (100ml/min^−1^), Fe CO_2_ is the concentration of CO_2_ exiting and Fi CO_2_ is the concentration CO_2_ entering the respirometer. Species were randomly assigned across the respiration chambers and the order in which flies were assayed in (chamber order) was corrected for statistically (see below).

### Body Size

To check for any potential effect of body size differences between species, wing length was measured as a proxy for body size [54]. A mean of 26 (range 20-30) males of each species were collected and immediately stored in ethanol during the collections for the viral load assay. Subsequently, wings were removed and photographed under a dissecting microscope. Using ImageJ software (version 1.48) the length of the IV longitudinal vein from the tip of the proximal segment to where the distal segment joins vein V was recorded, and the mean taken for each species.

### Host phylogeny

The host phylogeny was inferred as described in Longdon *et al* (2015) [35], using the *28S, Adh, Amyrel, COI, COII, RpL32* and *SOD* genes. Briefly, any publicly available sequences were downloaded from Genbank, and any not available we attempted to sanger sequence [34]. In total we had *RpL32* sequences for all 45 species, 41 for *28s*, 43 for *Adh*, 29 for *Amyrel*, 38 for *COI*, 43 for *COII* and 25 for *SOD* (see Figshare 10.6084/m9.figshare.6653192 for full details). The sequences of each gene were aligned in Geneious (version 9.1.8, [55]) using the global alignment setting, with free end gaps and a cost matrix of 70% similarity. The phylogeny was constructed using the BEAST program (version 1.8.4,[56]). Genes were partitioned into three groups each with their own molecular clock models. The three partitions were: mitochondrial *(COI, COII);* ribosomal (28S); and nuclear *(Adh, SOD, Amyrel, RpL32).* A random starting tree was used, with a relaxed uncorrelated lognormal molecular clock. Each of the partitions used a HKY substitution model with a gamma distribution of rate variation with 4 categories and estimated base frequencies. Additionally, the mitochondrial and nuclear data sets were partitioned into codon positions 1+2 and 3, with unlinked substitution rates and base frequencies across codon positions. The tree-shape prior was set to a birth-death process. The BEAST analysis was run twice to ensure convergence for 1000 million MCMC generations sampled every 10000 steps. The MCMC process was examined using the program Tracer (version 1.6, [57]) to ensure convergence and adequate sampling, and the constructed tree was then visualised using FigTree (version 1.4.3, [58]).

### Statistical analysis

All data were analysed using phylogenetic mixed models to look at the effects of host relatedness on viral load across temperature. We fitted all models using a Bayesian approach in the R package MCMCglmm [59,60]. We ran trivariate models with viral load at each of the three temperatures as the response variable similar to that outlined in Longdon *et al.* (2011) [34]. The models took the form:

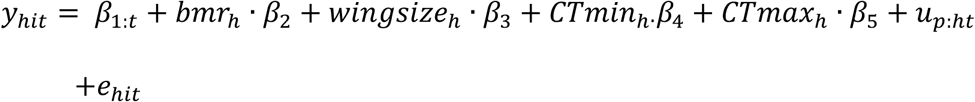

Where *y* is the change in viral load of the *i*^*th*^ biological replicate of host species *h*, for temperature *t* (high, medium or low). *β* are the fixed effects, with *β*_*1*_ being the intercepts for each temperature, *β2* being the effect of basal metabolic rate, *β3* the effect of wing size, and *β4* and *β5* the effects of the critical thermal maximum (CTmax) and minimum (CTmin) respectively. *u_p_* are the random phylogenetic species effects and *e* the model residuals. We also ran models that included a non-phylogenetic random species effect (*u_np:ht_*) to allow us to estimate the proportion of variation explained by the host phylogeny [34,35,61]. We do not use this term in the main model as we struggled to separate the phylogenetic and non-phylogenetic terms. Our main model therefore assumes a Brownian motion model of evolution [62].

The random effects and the residuals are assumed to be multivariate normal with a zero mean and a covariance structure **V_p_** ⊗ **A** for the phylogenetic affects and **Ve** ⊗ **I** for the residuals. **A** is the phylogenetic relatedness matrix, **I** is an identity matrix and the **V** are 3×3 (co)variance matrices describing the (co)variances between viral titre at different temperatures. The phylogenetic covariance matrix, **V_p_**, describes the inter-specific variances in each trait and the inter-specific covariances between them. The residual covariance matrix, **V_e_**, describes the within-species variance that can be both due to real within-species effects and measurement or experimental errors. The off-diagonal elements of **V_e_** (the covariances) are not estimable because no vial has been subject to multiple temperatures and so were set to zero. We excluded *D. pseudoobscura* from the full model as data for BMR was not collected, but included it in models that did not include any fixed effects, which gave equivalent results.

Diffuse independent normal priors were placed on the fixed effects (means of zero and variances of 10^8^). Parameter expanded priors were placed on the covariance matrices resulting in scaled multivariate F distributions which have the property that the marginal distributions for the variances are scaled (by 1000) F _1,1_. The exceptions were the residual variances for which an inverse-gamma prior was used with shape and scale equal to 0.001. The MCMC chain was run for 130 million iterations with a burn-in of 30 million iterations and a thinning interval of 100,000. We confirmed the results were not sensitive to the choice of prior by also fitting models with inverse-Wishart and flat priors for the variance covariance matrices (described in [34]), which gave qualitatively similar results (data not shown). All confidence intervals (CI’s) reported are 95% highest posterior density intervals.

Using similar model structures we also ran a univariate model with BMR and a bivariate model with CT_min_ and CT_max_ as the response variables to calculate how much of the variation in these traits was explained by the host phylogeny. Both of these models were also run with wing as a proxy for body size as this is known to influence thermal measures [51]. We observed significant levels of measurement error in the metabolic rate data; this was partially caused by respiratory chamber order during the assay. We corrected for this in two different ways. First, we fitted a linear model to the data to control for the effect of respiratory chamber number and then used this corrected data in all further models. We also used a measurement error model that controls for both respiratory chamber number effects and random error. Both of these models gave similar results although the measurement error model showed broad CIs suggesting the BMR data should be interpreted with caution. All datasets and R scripts with the model parameterisation are provided as supplementary materials.

## Results

To investigate the effect of temperature on virus host shifts we quantified viral load in 12,827 flies from 45 species of Drosophilidae at three temperatures (Fig 1). DCV replicated in all host species, but viral load differed between species and temperatures (Fig 1). Species with similar viral loads cluster together on the phylogeny (Fig 2). Measurements were highly repeatable (Table 1), with a large proportion of the variance being explained by the inter-specific phylogenetic component *(v*_*p*_), with little within species or measurement error *(v*_*r*_) (Repeatability= *v_p_/(y_p_ + v_r_):* Low = 0.90 (95% CI: 0.84, 0.95), Medium = 0.96 (95% CI: 0.93, 0.98), and High = 0.95, (95% CI: 0.89, 0.98)). We also calculated the proportion of between species variance that can be explained by the phylogeny as *vp/(vp+* vs) [63], which is equivalent to Pagel’s lambda or phylogenetic heritability [61,64]. We found the host phylogeny explains a large proportion of the interspecific variation in viral load across all three temperatures, although these estimates have broad confidence intervals due to the model struggling to separate the phylogenetic and non-phylogenetic components (Low = 0.77, 95% CI: 0.28, 0.99; Medium = 0.53, 95% Cl: 0.31×l0^−5^, 0.85; High = 0.40, 95% Cl: 0.99×l0^−5^, 0.74).

**Fig 1.**
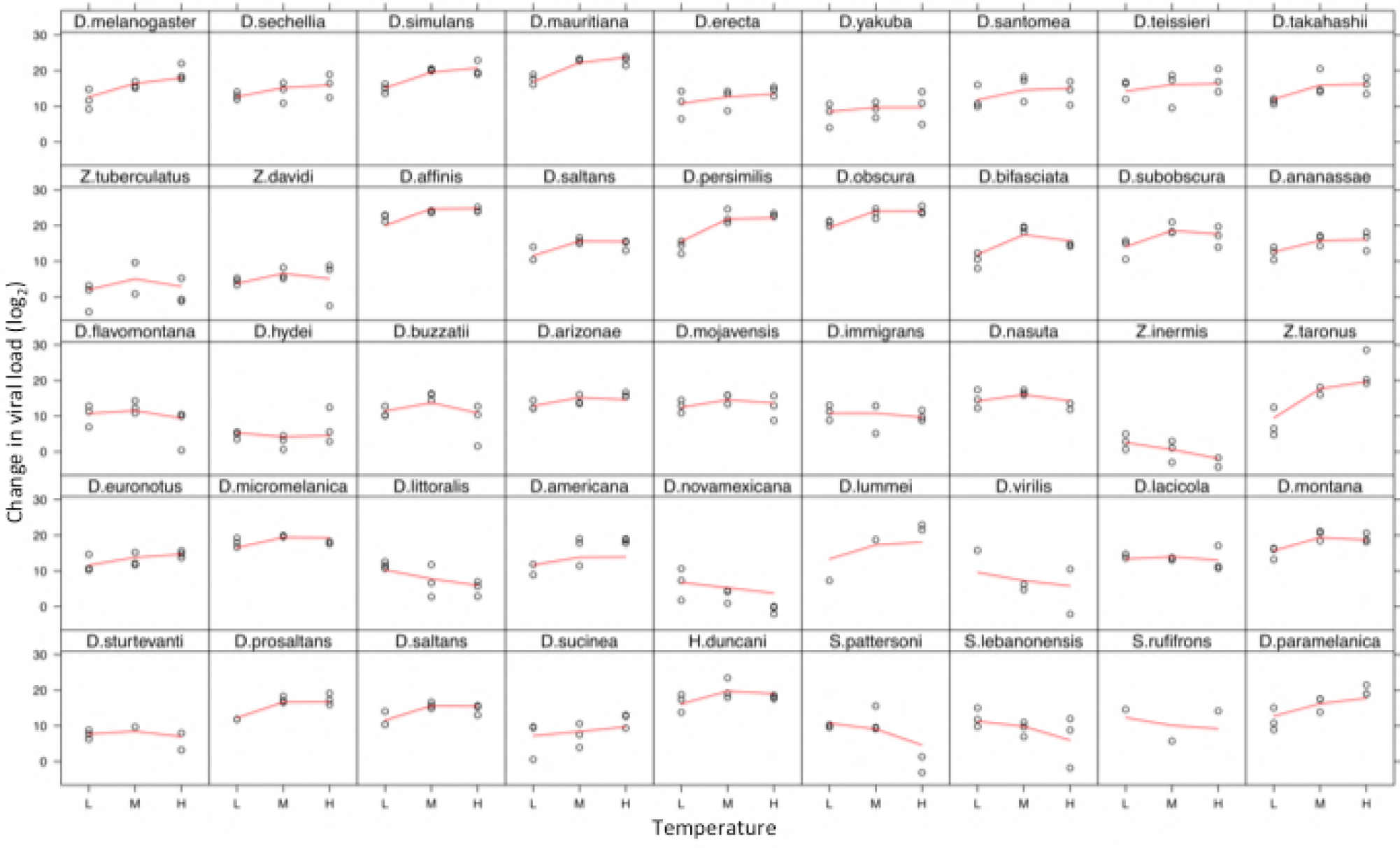
Change in viral load (log_2_) for 45 *Drosophilidae* species across three temperatures ( Low= 17°C, Medium=22°C and High=27°C). Individual points are for each replicate (change in viral load between day 0 and day 2 post infection), the red line is the predicted values from the phylogenetic mixed model. Panels are ordered by the tips on the phylogeny as in Fig 2.

**Fig 2.**
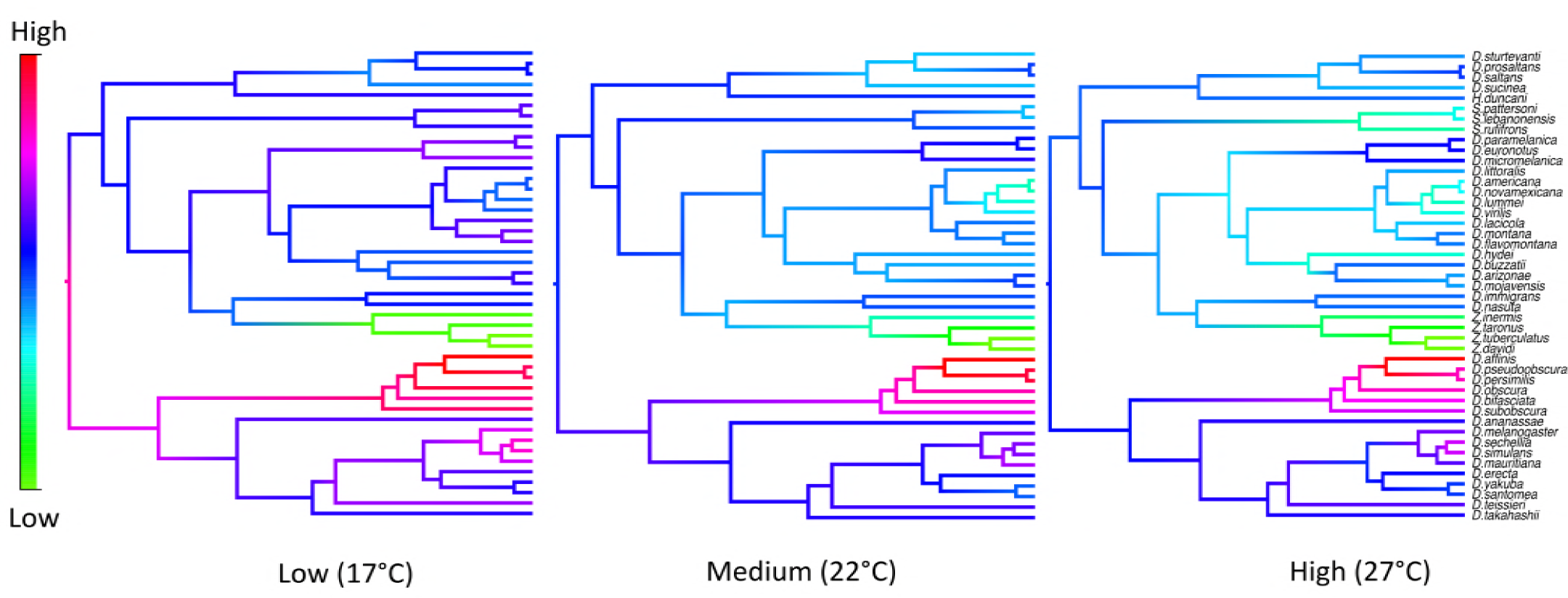
Ancestral state reconstructions to visualise the change in viral load across the host phylogeny at three temperatures. Ancestral states are plotted as colour gradients across the tree. The colour gradient represents the change in RNA viral load; red represents the highest and green the lowest viral load at that temperature. Ancestral states were estimated using a phylogenetic mixed model that partitioned the inter-specific variance into that explained by the host phylogeny under a Brownian model of evolution (*v*_*p*_), and a species-specific variance component that is not explained by the phylogeny (*v*_*s*_).

**Table 1.** Change in viral load with temperature.

| Temperature | Intercepts |  | Between-species |  | Within-species |  |
| --- | --- | --- | --- | --- | --- | --- |
| | | | Variance ($v_p$) | | Variance ($v_r$) | |
|  | Mean | 95% CIs | Mean | 95% CIs | Mean | 95% CIs |
| **Low** | 11.9 | 9.5, 14.6 | 65.3 | 32.3, 110.3 | 6.9 | 4.8, 9.3 |
| **Medium** | 14.3 | 11.7, 17.1 | 172.2 | 90.2, 278.8 | 7.0 | 4.8, 9.2 |
| **High** | 13.5 | 10.8, 16.7 | 260.6 | 119.7, 413.7 | 12.8 | 8.9, 17.5 |

Intercepts are the temperature-specific intercepts when the other covariates (e.g. wing size) are set to their temperature specific means. They can be interpreted as the expected viral loads at the root of the phylogeny at each temperature. $v_p$ is the variance in between-species effects, which are structured by the phylogeny, and $v_r$ is the variance in within species effects attributable to between individual differences and measurement error.

To examine if species responded in the same or different way to changes in temperature we examined the relationships between susceptibilities across the different temperatures. We found strong positive phylogenetic correlations between viral loads across the three temperatures (Table 2). Our models showed that the variance in viral load increased with temperature, whilst the mean viral load showed no such upward trend (Table 1), suggesting that the variance changes are not just due to scaling effects.

**Table 2.** Interspecific correlations between viral loads at each temperature.

| Temperatures | Interspecific Correlation | 95% CIs |
| --- | --- | --- |
| **High-Low** | 0.89 | 0.77, 0.98 |
| **Medium-Low** | 0.92 | 0.90, 0.99 |
| **Medium-High** | 0.97 | 0.93, 0.99 |

The high correlations suggest the rank order of susceptibility of the species is not changing with increasing temperature. However, the change in variance suggests that although the reaction norms are not crossing they are diverging from each other as temperature increases i.e. the most susceptible species are becoming more susceptible with increasing temperature, and the least susceptible less so [65]. For example, *D. obscura* and *D. affinis* are the most susceptible species at all three temperatures. The responses of individual species show that some species have increasing viral load as temperature increases (Fig 1, e.g. *Z. taronus, D. lummei)*, whilst others decease (e.g. *D. littoralis, D. novamexicana).*

The changes we observe could be explained by the increase in temperature effectively increasing the rate at which successful infection is progressing (i.e. altering where in the course of infection we have sampled). However, this seems unlikely as at 2 days post infection at the medium temperature (22°C), viral load peaks and then plateaus [35]. Therefore, in those species where viral load increases at higher temperatures the peak viral load itself must be increasing, rather than us effectively sampling the same growth curve but at a later time point. Likewise, in those species where viral load decreased at higher temperatures, viral load would need to first increase and then decrease, which we do not observe in a time course at 22°C [35]. To check whether this also holds at higher temperatures we carried out a time course of infection in a subset of six of the experimental species at 27°C, where we would expect the fastest transition between the rapid viral growth and the plateau phase of infection to occur (S1 Fig), This allowed us to confirm that the decreasing viral loads observed in some species at higher temperatures are not due to general trend for viral loads to decline over longer periods of (metabolic) time.

We quantified the lower and upper thermal tolerances (CT_min_ and CT_max_) across all 45 species with 3 replicates per species. Neither CT_max_ nor CT_min_ were found to be significant predictors of viral load (CT_min_ −0.21, 95% CI: −0.79, 0.93, pMCMC = 0.95 and CT_max_ 0.31, 95% CI: −0.11, 0.74, pMCMC = 0.152). When treated as a response in models we found the host phylogeny explained a large proportion of the variation in thermal maximum (CT_max_: 0.95, 95% CI: 0.84, 1) and thermal minima (CT_min_: 0.98, 95% CI: 0.92, 0.99, S2 Fig).

We also measured the basal metabolic rate of 1320 flies from 44 species, across the three experimental temperatures, to examine how cellular function changes with temperature. BMR was not found to be a significant predictor of viral load when included as a fixed effect in our model (slope = 9.09, 95% CI = -10.13, 20.2689, pMCMC = 0.548).

BMR increased with temperature across all species (mean BMR and SE: Low 0.64 ± 0. 02, Medium 1.00 ± 0.04, High 1.2 ± 0.04 CO_2_ml/min^−1^, S3 Fig).

When BMR was analysed as the response in models, the phylogeny explained a small amount of the between species variation (Low 0.19, 95% CI: 2 × 10^−8^, 0.55, Medium 0.10, 95% CI: 5 × 10^−7^, 0.27, High 0.03, 95% CI: 8 × 10^−9^ - 0.13, S4 Fig) indicating high within species variation or large measurement error. Consequently the species/temperature mean BMRs used in the analysis of viral load will be poorly estimated and so the effects of BMR will be underestimated with too narrow credible intervals. To rectify this we ran a series of measurement error models, the most conservative of which gave a slope of -9.8 but with very wide credible intervals (−62.5, 42.6). Full details of these models are given in the Supplementary Materials.

## Discussion

We found that susceptibilities of different species responded in different ways to changes in temperature. The susceptibilities of different species showed either increases or decreases at higher temperatures. There was a strong phylogenetic correlation in viral load across the three experimental temperatures (Table 2). However, the variance in viral load increased with temperature, whereas the mean viral load did not show the same trend. This suggests that the rank order of susceptibility of the species remains relatively constant across temperatures, but as temperature increases the most susceptible species become more susceptible, and the least susceptible less so.

Changes in global temperatures are widely predicted to alter host-parasite interactions and therefore the likelihood of host shifts occurring [5,18,66–68]. The outcome of these interactions may be difficult to predict if temperature causes a different effect in the host and pathogen species [15,69–72]. Our results show that changes in temperature may change the likelihood of pathogens jumping into certain species, although they suggest that it may not alter which species are the most susceptible to a novel pathogen.

The increase in phylogenetic variance with temperature is effectively a form of genotype-by-environment interaction [25,73–75]. However, it varies from the classically considered ecological crossing of reaction norms, as we do not see a change in the rank order of species susceptibly. Instead, we find the species means diverge with increasing temperatures and so the between species differences increase [65,76].

As temperature is an important abiotic factor in many cellular and physiological processes, we went on to examine the underlying basis of why viral load might change with temperature. Previous studies that found phylogenetic signal in host susceptibility were carried out at a single experimental temperature [34,35]. Therefore, the patterns observed could potentially be explained by some host clades being assayed at sub-optimal thermal conditions. We used CT_max_ and CT_min_ as proxies for thermal optima, which due to its multifaceted nature is problematic to measure directly [77–79]. We also measured basal metabolic rate across three temperatures to see if the changes in viral load could be explained by general increases in enzymatic processes. We found that these measures were not significant predictors of the change in viral load with temperature.

The host immune response and cellular components utilised by the virus are likely to function most efficiently at the thermal optima of a species, and several studies have demonstrated the outcomes of host-pathogen interactions can depend on temperature [23,25,69,75]. However, the mechanisms underlying the changes in susceptibility with temperature seen in this study are uncertain and a matter for speculation. Our results show that in the most susceptible species, viral load increases with temperature; this may be due to the virus being able to successful infect and then freely proliferate, utilizing the host cells whist avoiding host immune defences. In less susceptible species viral load does not increase with temperature, and in some cases it actually appears to decreases. Here, temperature may be driving an increase in biological processes such as enhanced host immunity, or simply increasing the rate of degradation or clearance of virus particles that have failed to establish an infection of host cells.

In conclusion, we have found changes in temperature can both increase or decrease the likelihood of a host shift. Our results show the rank order of species susceptibilities remain the same across temperatures, suggesting that studies of host shifts at a single temperature can be informative in predicting which species are the most vulnerable to a novel pathogen. Understanding how environmental factors might affect broader taxonomic groups of hosts and pathogens requires further study if we are to better understand host shifts in relation to climate change in nature.

## Acknowledgements

Many thanks to Darren Obbard and Frank Jiggins for useful discussion and Vanessa Kellerman and Johannes Overgaard for discussion about thermal assay methods. Thanks to Dave Hosken for use of BMR chambers and to the Drosophila Species Stock Centre for supplying flies.

## Supporting information

**Supplementary methods:** Set up and equipment for carrying out the CTmax/CTmin assays. Measurement correction model methods

**S1 Table:** Full list of species used in the experiment and their rearing food for stock populations.

**S2 Table:** Genbank accession numbers of sequences used to infer the host phylogeny.

**S1 Fig:** a) Viral load for 6 *Drosophilidae* species at 27°C, on day 0, day 1 and day 2 post infection. b) Change in viral load for the same 6 *Drosophilidae* species across the three temperatures

**S2 Fig:** Ancestral state reconstructions of CTmin and CTmax for experimental flies.

**S3 Fig:** Change in Basal Metabolic Rate (BMR) for 44 *Drosophilidae* species across three temperatures.

**S4 Fig:** Ancestral state reconstructions of BMR across the three experimental temperatures

